# Distinct memory CD4 subset tropism of two CCR5-tropic HIV-1 in a rapid progressor

**DOI:** 10.1101/2025.01.21.634053

**Authors:** Manukumar Honnayakanahalli Marichannegowda, Yasmine Farah, Meera Bose, Eric Sanders-Buell, David King, Leilani Francisco, Leigh Anne Eller, Sodsai Tovanabutra, Nelson L. Michael, Merlin L. Robb, Hongshuo Song

## Abstract

CCR5 tropic HIV-1 mainly infect memory CD4^+^ T cells. Memory CD4^+^ T cells are heterogenous which comprise central memory (CM), transitional memory (TM) and effector memory (EM) subsets based on cell differentiation stages. Previous studies demonstrate that low viral burden in the CM subset correlates with long-term non-progressive HIV-1 infection (1, 2) and is a hallmark of non-pathogenetic SIV infection (3, 4). These findings indicate the importance of CD4 subset tropism of HIV/SIV in determining disease progression. However, an important, yet unanswered question is whether CCR5 HIV-1 variants have different memory CD4 subset preferences. Here, we demonstrate clear compartmentalization of two CCR5 HIV-1 in different memory CD4 subsets *in vivo*.

## Main text

Participant 40512 was identified in the RV217 cohort (5). Longitudinal virus sequencing demonstrated that 40512 was initially infected by a single transmitted/founder (T/F) virus and was superinfected on day 401 (Supplementary Figure 1). While 40512 did not harbor CXCR4 virus (6), he had the fastest CD4 decline among participants in the same cohort who harbored only CCR5 virus (0.76 cells/μL/day vs. 0.23 cells/μL/day) (Supplementary Figure 2A). The viral load (VL) set point in 40512 was relatively high (Supplementary Figure 2B). The rapid CD4 drop occurred before superinfection (Figure 1A), suggesting that it was caused by the T/F virus rather than superinfection. After superinfection, the superinfecting (SI) strain was predominant in plasma, while the original T/F became a minor lineage (Figure 1B and Supplementary Figure 1).

**Figure 1.**
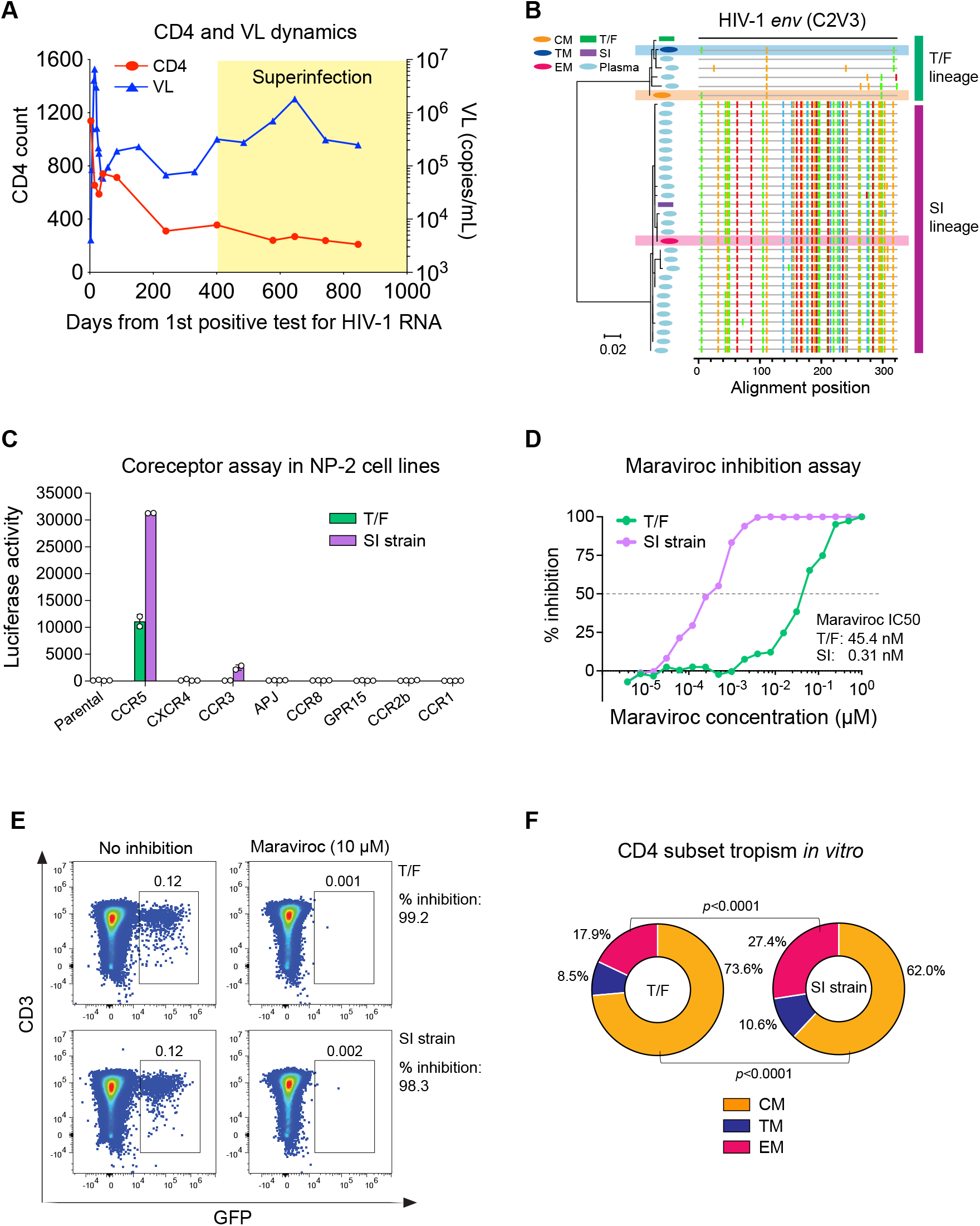
Identification and characterization of the T/F virus and the superinfecting strain. **(A)** CD4 and VL dynamics in 40512. **(B)** Phylogenetic tree and highlighter plot showing the relationship between viruses in plasma and in each CD4 subset. Different viruses are color-coded. **(C)** Coreceptor usage of the T/F virus and SI strain in NP-2 cell lines. **(D)** Sensitivity to Maraviroc. The Maraviroc IC_50_ of each virus is shown. **(E)** Sensitivity to Maraviroc in primary CD4^+^ T cells. **(F)** *In vitro* CD4 subset tropism. The percentage of each CD4 subset among the total infected cells is shown. The statistical significance is determined by a Chi-squared test.

Like other participants who harbored only CCR5 virus, cell-associated HIV-1 RNA was undetectable in the naïve CD4 subset in 40512 but was detected in all memory subsets (6). Sequencing of HIV-1 RNA in each memory subset identified compartmentalization of the T/F and the superinfecting strain (Figure 1B). While the T/F lineage was replicating in the CM and TM subsets, the superinfecting lineage was replicating in the EM subsets (Figure 1B). Phylogenetic analysis showed that the superinfecting strain, which was predominant in plasma, was originated from the EM subset, while the T/F lineage was originated from the CM and TM subsets (Figure 1B). These data demonstrate that the T/F virus and the superinfecting strain preferentially infect different memory CD4 subsets *in vivo*. The EM subset, which was infected by the superinfecting strain, released more virions into the plasma than the CM and TM subsets.

Coreceptor assay confirmed that both the T/F virus and the superinfecting strain are CCR5 tropic (Figure 1C). The infectivity of both viruses can be completely inhibited by 1 μM of the CCR5 inhibitor Maraviroc. However, the Maraviroc IC_50_ of the T/F virus was 146-fold higher than the superinfecting strain (45.5 nM vs. 0.31 nM) (Figure 1D). This data suggests the possibility that these two viruses might use CCR5 in different ways. In primary CD4^+^ T cells, the infectivity of both viruses can be nearly completely inhibited by 10 μM Maraviroc (Figure 1E). Therefore, both viruses rely on CCR5 to enter primary CD4^+^ T cells, while the contributions of other coreceptors are very minimal, if any.

We next determined whether these two viruses preferentially infect different CD4 subsets *in vitro* by using GFP pseudovirus. Among the cells infected by the T/F, 73.6% were CM, 8.5% were TM, and 17.9% were EM. Among the cells infected by the SI strain, 62% were CM, 10.6% were TM, and 27.4 % were EM (Figure 1F). This single-round infection shows that the T/F has an advantage to infect the CM subset than the SI strain (*p* < 0.0001), while the SI strain has an advantage to infect the EM subset than the T/F (*p* < 0.0001). Therefore, their compartmentalization *in vivo* could be at least in part determined at the entry level. It is possible that even a small difference in entry ability could be amplified after multiple rounds of replication, consequently leading to compartmentation in different CD4 subsets *in vivo*.

In summary, the current study demonstrates that CCR5 HIV-1 variants have different memory CD4 subset preferences. Several important questions remain. First, the mechanisms in determining the CD4 subset tropism of CCR5 HIV-1. Second, because the EM cells are mainly distributed in the mucosal sites where most HIV-1 transmissions occur, while the CM cells are mainly in the lymph nodes and are critical for CD4 homeostasis, the CD4 subset tropism in determining HIV-1 transmissibility and pathogenesis requires future investigation. Addressing these questions is expected to provide deeper insights into HIV-1 prevention and functional cure.

