## Supplementary file for "Distinct memory CD4 subset tropism of two CCR5-tropic HIV-1 in a rapid progressor"

### Supplementary information

#### Study participants

Participant 40512 was identified from the RV217 Thailand acute infection cohort (1). The RV217 cohort followed at risk participants twice weekly to identify HIV-1 RNA within a few days of the last HIV-1 RNA negative test, thus allowing investigation of HIV-1 evolution and host immune response from the earliest stage of HIV-1 infection. Both the transmitted/founder (T/F) virus and the superinfecting virus in 40512 were CRF01_AE. For CD4 and viral load analysis, we included 12 participants in the RV217 Thailand acute infection cohort who were infected by CCR5 tropic T/F virus and did not undergo coreceptor switch for the first two years of infection as we reported in a previous study (2). The study was approved by the local ethics review boards, the Walter Reed Army Institute of Research, and the institutional review boards of the University of Maryland School of Medicine (number: HP-00089353). Written consent was provided by all participants.

#### HIV-1 sequence analysis

Longitudinal envelope sequences of 40512 were retrieved from GenBank (MN792521-MN792549; OM826457-OM826666). To amplify the C2V3 region of the HIV-1 *env* in each CD4 subset, HIV-1 RNA was extracted using the RNeasy Mini kit (Qiagen). A total of 8.5 µL extracted RNA was subjected to reverse transcription. The cDNA was synthesized using the SuperScript III reverse transcriptase (Invitrogen) using the primer 1.R3.B3R 5’-ACTACTTGAAGCACTCAAGGCAAGCTTTATTG-3’ (nt 9642-9611 in HXB2). The first round PCR was carried out using the forward primer 5'-ACAGTRCARTGYACACATGG-3' (nt 6954-6973) and the reverse primer 5'-CACTTCTCCAATTGTCCITCA-3' (nt 7648-7668). The first round PCR conditions were as follow: one cycle at 94°C for 2 min; 30 cycles of a denaturing step at 94°C for 15 sec, an annealing step at 55°C for 30 sec, an extension step at 68°C for 1 min; and one cycle of an additional extension at 68°C for 10 min. A total of 5 µL of first round PCR products were used for second round PCR amplification. The second round PCR was carried out using the forward primers 5'-CAGTACAATGYACACATGG-3' (nt 6955-6973) and reverse primer 5'-AGAAAAATTCYCCTCYACAA-3' (nt 7374-7355). The second round PCR conditions were as follow: one cycle at 94°C for 2 min; 35 cycles of a denaturing step at 94°C for 15 sec, an annealing step at 55°C for 30 sec, an extension step at 68°C for 1 min; and one cycle of an additional extension at 68°C for 10 min. All PCR amplifications were carried out using the Platinum™ Taq DNA Polymerase High Fidelity (Invitrogen). The PCR amplicons were directly sequenced by the cycle sequencing and dye terminator methods. Individual sequences were assembled and edited using Sequencher software (Gene Codes). The sequences were aligned using the Gene Cutter tool (https://www.hiv.lanl.gov/content/sequence/GENE_CUTTER/cutter.html) in the Los Alamos HIV Sequence Database, followed by manual adjustment to obtain the optimal alignment. The phylogenetic tree was constructed by MEGA6 using the maximum likelihood method (3). The Highlighter plot was generated using the Highlighter tool in the Los Alamos HIV Sequence Database (https://www.hiv.lanl.gov/content/sequence/HIGHLIGHT/highlighter_top.html).

**Pseudovirus preparation and titration**

The full-length *env* sequences of the 40512 T/F virus (defined as the day 6 consensus sequence) and the superinfecting strain (defined as the consensus sequence of the superinfecting lineage at day 401) were chemically synthesized (GenScript, USA) and cloned into the expression vector pcDNA3.3-TOPO (Invitrogen, USA). Pseudovirus stocks were prepared as previously described (4). In brief, 2 µg of HIV-1 *env* clone was co-transfected with 4 µg of the pNL4.3-ΔEnv-vpr+-luc+ backbone (for coreceptor assay in NP-2 cell lines) (4), or 4 µg of the pNL4.3-ΔEnv EGFP backbone (5) (for HIV-1 entry assay in primary $CD4^+$ T cells) into 293T cells using the FuGENE6 transfection reagent (Promega, USA). The cells were cultured at 37°C for 6 hours, after which the culture medium was completely replaced by fresh medium. The culture supernatants were harvested at 72 hours post transfection, aliquoted and stored at −80°C until use. The infectious titers ($TCID_{50}$) of the pseudovirus stocks were titrated on TZM-bl cells (obtained from the HIV Reagent Program, ARP-8129).

**Coreceptor usage assay**

HIV-1 coreceptor usage was determined in a panel of NP-2 cell lines expressing CD4 together with different G protein-coupled receptors (CCR5, CXCR4, APJ, CCR3, CCR8, CCR1 and CCR2b) as previously described (4). The parental NP-2 cell line expressing CD4 alone was used as a control. In brief, NP-2 cell lines expressing different coreceptors were seeded into a 96-well plate at a density of $1 \times 10^5$ cells per well one day before infection. The next day, the cells were infected with approximately 200 $TCID_{50}$ of each pseudovirus (MOI = 0.002) containing the luciferase reporter. After 6 hours of infection at 37°C, the cells were washed three times and cultured at 37°C for three days. Three days after infection, the cells were lysed, and the infectivity in each well was determined by measuring the RLUs in the cell lysates using the Britelite plus system (PerkinElmer, USA). Viral infection was considered positive if the RLU value was more than 5-fold higher than the background RLU value in the parental cell line.

**Maraviroc inhibition assay in NP-2 CCR5 cell line**

To determine virus sensitivity to the CCR5 inhibitor Maraviroc, NP-2 CCR5 cells were seeded in a 96-well plate at a density of $1 \times 10^5$ cells per well one day before infection. The next day, the cells were pre-treated with 1:2 serial diluted Maraviroc (starting from 1.0 μM) at 37°C for 1 hour. The Maraviroc-treated cells were then infected with approximately 500 $TICD_{50}$ of each pseudovirus containing the luciferase report and cultured at 37°C for three days. Three days after infection, the infected cells were lysed. The infectivity in each well was determined by measuring the RLU in the cell lysate using the Britelite plus system (PerkinElmer). The percentage of Maraviroc inhibition in each well was determined by comparing the RLU values with the positive control wells without Maraviroc inhibition. The $IC_{50}$ of Maraviroc was determined using a linear regression model.

**Determination of virus infectivity and sensitivity to Maraviroc in primary $CD4^+$ T cells**

Primary $CD4^+$ T cells from a healthy donor were purified from PBMCs by negative selection (Miltenyi Biotec, USA). Purified $CD4^+$ T cells were stimulated for three days by 1 µg/mL soluble anti-CD3 (clone OKT3, eBioscience, USA) and 1 µg/mL soluble anti-CD28 (clone CD28.2, eBioscience, USA) in the presence of 50 IU/mL IL-2 (PeproTech, USA). One million of stimulated $CD4^+$ T cells were infected over night by each pseudovirus containing the GFP reporter (MOI = 0.1). The infected cells were washed three times with RPMI1640 after infection and were cultured in a 24-well plate with RPMI1640 containing 10% FBS and 50 IU/mL IL-2. Three days after infection, the percentage of GFP positive cells were determined by flow cytometry. To determine virus sensitivity to Maraviroc in primary $CD4^+$ T cells, one million stimulated $CD4^+$ T cells were pre-treated with 10 µM of Maraviroc at 37°C for 1 hour before infection. The infection was carried out as described above. The percentage of GFP positive cells were determined by flow cytometry three days after infection.

**CD4 subset analysis and sorting**

PBMCs from participant 40512 were stained by the following antibodies: CD3-Brilliant Violet 605 (clone OKT3, BioLegend), CD4-PerCP-Cy5.5 (clone OKT4, BioLegend), CCR7-PE-CF594 (clone 2-L1-A, BD Biosciences), CD27-PE (clone M-T271, BioLegend), CD45RO-APC (clone UCHL1, BioLegend). The cells were then stained by the Live-dead aqua prior to flow analysis (Invitrogen). The stained cells were sorted on a BD FACSAria II cell sorter (BD Biosciences). Four CD4 subsets were defined as follows: naïve ($CD45RO^-$, $CCR7^+$, and $CD27^+$), central memory ($CD45RO^+$, $CCR7^+$, and $CD27^+$), transitional memory ($CD45RO^+$, $CCR7^-$, and $CD27^+$) and effector memory

($CD45RO^+$, $CCR7^-$, and $CD27^-$). The purity of each sorted subset was higher than 95%. All commercial antibodies were validated by the vendors.

**Analysis of virus CD4 subset tropism *in vitro***

Primary $CD4^+$ T cells from a healthy donor were purified from PBMCs by negative selection (Miltenyi Biotec, USA). Purified $CD4^+$ T cells were stimulated for three days by 1 μg/mL soluble anti-CD3 (clone OKT3, eBioscience, USA) and 1 μg/mL soluble anti-CD28 (clone CD28.2, eBioscience, USA) in the presence of 50 IU/mL IL-2 (PeproTech, USA). One million of stimulated $CD4^+$ T cells were infected over night by pseudovirus containing the GFP reporter (MOI = 0.1). Three days after infection, the cells were stained by the following antibodies: CD3-eFluor 450 (clone OKT3, eBioscience), CD4-Pe-Cy7 (clone OKT4, Biolegend), CCR7-PE-CF594 (clone 2-L1-A, BD Biosciences), CD27-PE (clone M-T271, BioLegend), and CD45RO-APC (clone UCHL1, BioLegend). The infected cells were then analyzed by flow cytometry. The phenotype of the GFP positive cells were determined as follows: naïve ($CD45RO^-$, $CCR7^+$, and $CD27^+$), central memory ($CD45RO^+$, $CCR7^+$, and $CD27^+$), transitional memory ($CD45RO^+$, $CCR7^-$, and $CD27^+$) and effector memory ($CD45RO^+$, $CCR7^-$, and $CD27^-$). The proportion of each CD4 subset among the total GFP positive cells was calculated. All commercial antibodies were validated by the vendors.

**Statistical analysis**

The difference between the percentage of each infected CD4 subset was compared using a Chi-squared test. The rate of CD4 decline was determined using a linear mixed effect model (LME) as we previously described (2). The LME model was hierarchical in the sense that it estimated a population specific slope and intercept with time, as well as subject-specific slopes and intercepts. Longitudinal CD4 data from the earliest available time point to the last available time point before ART initiation was used for the analysis.

**Data availability statement**

HIV-1 sequences used in the current study were previously deposited in GenBank (MN792521-MN792549; OM826457-OM826666). All data will be available upon request through the corresponding author.

**Acknowledgements**

The authors thank the study participants of the RV217 Thailand cohort. We thank the flow cytometry core of the University of Maryland School of Medicine for technical assistance. This study was supported by the Institute of Human Virology, University of Maryland School of Medicine and the NIH grant R01AI181601. Part of the study was supported by cooperative agreements between the Henry M. Jackson Foundation for the Advancement of Military Medicine, Inc., and the U.S. Department of Defense (DOD).

### Supplementary Figures

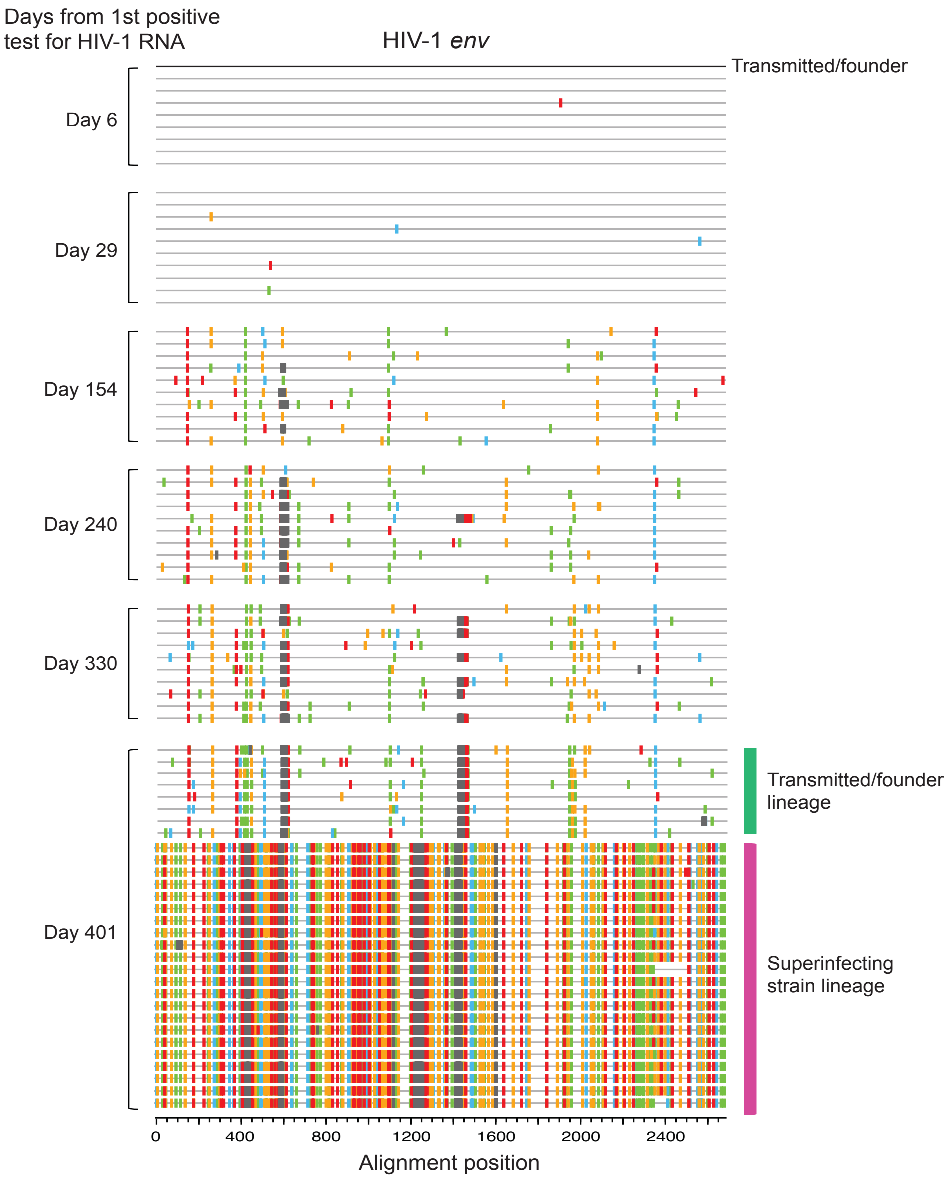


**Supplementary Figure 1. Highlighter plot showing longitudinal HIV-1 *env* sequences from participant 40512.** The T/F virus (defined as the consensus sequence of day 6) was used as the reference sequence. The superinfecting strain was firstly detected at day 401. The T/F lineage and the superinfecting lineage at day 401 were color coded.

**A**

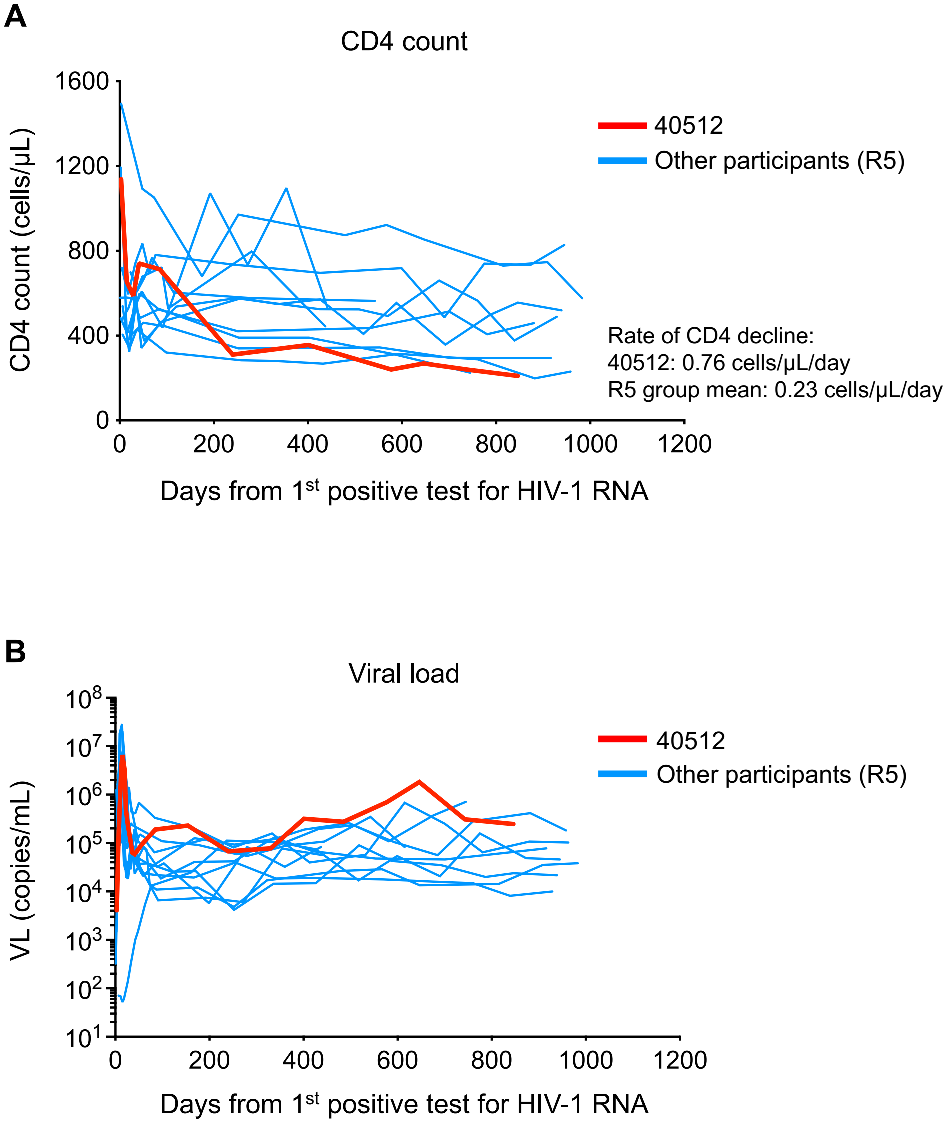


**Supplementary Figure 2. CD4 and VL dynamics in participant 40512 in comparison to other RV217 Thailand participants who harbored only CCR5 tropic HIV-1. (A)** CD4 dynamics in participant 40512 (red) in comparison to other RV217 Thailand participants who were infected by CCR5 tropic T/F virus and did not undergo coreceptor switch (blue). The rate of CD4 decline in 40512 and the mean of the entire group are shown. **(B)** VL dynamics in participant 40512 (red) in comparison to other RV217 Thailand participants who harbored only CCR5 tropic HIV-1 (blue).
